# mmSIM: An open toolbox for accessible structured illumination microscopy

**DOI:** 10.1101/2021.01.18.427184

**Authors:** Craig T. Russell, Michael Shaw

## Abstract

Since the first practical super-resolution structured illumination fluorescence microscopes (SIM) were demonstrated more than two decades ago the method has become increasingly popular for a wide range of bioimaging applications. The high cost and relative inflexibility of commercial systems, coupled with the conceptual simplicity of the approach and the desire to exploit and customise existing hardware, have led to the development of a large number of home-built systems. Several detailed hardware designs are available in the scientific literature, complemented by open-source software tools for SIM image validation and reconstruction. However, there remains a lack of simple open-source software to control these systems and manage the synchronization between hardware components, which is critical for effective SIM imaging. This article describes a new suite of software tools based on the popular Micro-Manager package, which enable the keen microscopist to develop and run a SIM system. We use the software to control two custom-built, high-speed, spatial light modulator-based SIM systems, evaluating their performance by imaging a range of fluorescent samples. By simplifying the process of SIM hardware development, we aim to support wider adoption of the technique.

## Introduction

Structured illumination microscopy (SIM) is a popular widefield super-resolution fluorescence microscopy technique which exploits frequency mixing between a spatially modulated illumination pattern and a sample fluorophore distribution to improve spatial resolution by a factor of up to two ^1,2^. The increased spatial resolution, coupled with the overall high quality of reconstructed images and fast acquisition speeds, have led to SIM being used in a wide range of bioimaging applications from the measurement of protein fibrillogenesis ^3,4^ to structural imaging of cells ^5^, organisms ^6^ and tissues ^7,8^. SIM also exposes samples to lower irradiance levels than other high and super-resolution resolution microscopy modalities, making it well suited for imaging of sensitive biological samples. The adoption of the method has also been aided by its’ innate compatibility with conventional fluorescent dyes and labels.

In the simplest case SIM relies on the capture of nine diffraction-limited images acquired under illumination of the sample with a series of sinusoidal excitation patterns. A single high-resolution image is then computationally reconstructed by inverting the forward frequency-mixing process ^1,9^. Alongside the conventional (diffraction-limited, zero order) object spectrum filtered by the optical transfer function (OTF) of the microscope objective, sinusoidal illumination, 1 + sin(***k***_1_***x***), downshifts additional high frequency information into the OTF support. The information contained within these (first order) frequency passbands can be computationally unmixed using three images captured as the phase of the illumination pattern is shifted. Knowledge of the wavevector (***k*_1_**) of the illumination pattern allows this high frequency information to be shifted back to its’ true location in Fourier space before the passbands are combined to yield an image with an extended frequency support (see Fig. 1). The separation of the zero and first order passbands also means that SIM image data can also be processed, by attenuation of information close to the passband centres, to remove out of focus light allowing fast, widefield, optically sectioned imaging ^10^. Alternatively, modifying the illumination pattern to introduce axial modulation allows this concept to be extended to 3D ^9^.

**Fig. 1.**
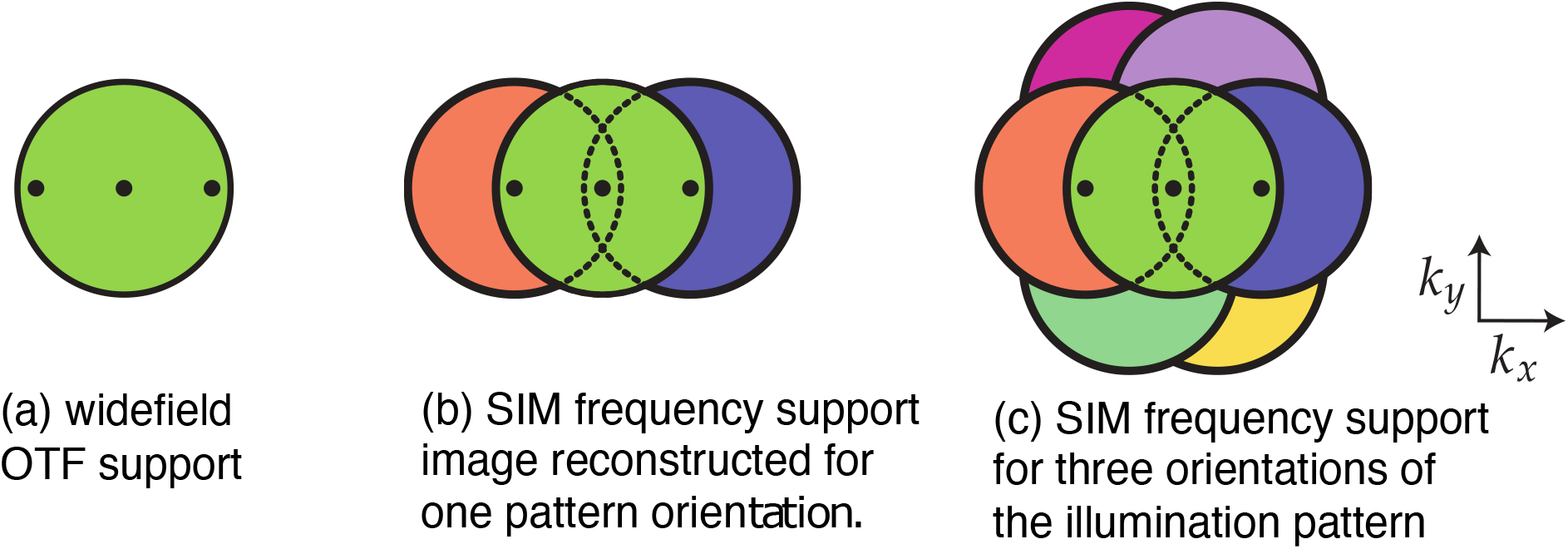
SIM reconstruction in Fourier space. (a) The circular area represents the diffraction limited OTF support (passband) of the objective lens. (b) Illumination of the sample with a sinusoidal irradiance pattern creates two additional passbands (blue and orange) which carry high spatial frequency information. SIM image reconstruction involves separating these passbands and shifting the information back to its origin in Fourier space. (c) Repeating this process using two further sinusoidal illumination patterns, oriented at 120° to the first pattern, isotropically extends the frequency support.

Early SIM systems typically relied on a diffraction grating inserted into the illumination light path of the microscope to generate a sinusoidal irradiance pattern ^1^. The phase and orientation of the illumination pattern was then controlled via mechanical translation and rotation of the grating. Most modern home-built SIM systems now use a spatial light modulator (SLM) such as liquid crystal on silicon (LCoS) ^11^ or digital micromirror device for pattern generation ^12^. In addition to a high imaging speed (refresh rates for these devices are typically on the order of kHz) and increased experimental robustness, these devices are extremely flexible allowing simple adjustment of the illumination pattern properties ^1,3^. A demagnified image of the SLM, usually illuminated with collimated laser light, is projected into the sample using off-the-shelf optical components. The SLM is programmed to display a series of binary phase gratings (bitplanes), and a spatial filter in the Fourier plane of the SLM removes unwanted diffracted orders. By synchronising the display of bitplanes with the exposure of the camera sensor, this configuration allows SIM images to be captured at many frames per second.

The wider adoption of SIM is limited both by the relatively high cost of commercial instruments as well as the time and expertise required to develop a self-built system. In the latter case, numerous SIM hardware designs exist in the literature and there are open source software tools for checking and validating raw SIM images ^14^ and reconstructing super-resolution images ^15^. However, there is a lack of open-source software tools to control SIM system hardware presenting a significant barrier to self-built SIM for the non-expert developer. The fairSIM-VIGOR ^16^ software project allows real time graphics processing unit (GPU)-enabled SIM image reconstruction using a hardware configuration based on multiple (acquisition) computers which pipe camera data to a separate reconstruction computer. However, the software requires its own bespoke frontend and an involved set-up process using a micro-controller to manage hardware synchronization and has limited compatibility with different hardware devices and customized setups.

In this article we describe an approach to SIM hardware control and image acquisition (mmSIM) based on the Micro-Manager (µManager)^17^ microscope control package which includes custom plugins for acquisition of SIM image sequences and real time visualization of the image power spectrum. Combined with a simple hardware design and a recipe for hardware synchronization, mmSIM enables the enthusiastic instrumentalist to build an affordable, robust, performant SIM system, that is easy to use and maintain.

## SIM hardware

### System design

We built a SIM system by adapting a commercial inverted microscope (IX71, Olympus) (see Fig. 2). Four diode lasers (405 nm-120 mW, 488 nm – 100 mW, 515 nm −100 mW, and 638 nm – 100mW) and a diode pumped solid state (DPSS) laser (561 nm-106 mW) (Luxx and Jive from Omicron-Laserage) controlled with an acousto-optic modulator were coupled into a single mode polarization maintaining fibre. At the fibre output laser light was collimated using a parabolic mirror to give a beam diameter of 1 mm. Optionally the beam was then passed through a Pockels cell (M350-80, Conoptics Inc.) and an achromatic quarter waveplate to allow fast control of the axis of linear polarisation. Corotating the polarization vector with the orientation of the illumination pattern ensures optimal pattern contrast ^18^, however we find it is only strictly necessary at high pattern wavevectors, i.e. when the first order diffracted beams from the SLM are incident close to the pupil edge of a high numerical aperture (NA) objective lens. Setting the direction of linear polarisation at this point in the optical system removes the need for additional beam expansion systems (due to the small aperture of the Pockels cell), but means that light reflected from the SLM is slightly elliptically polarised ^21^. However, this has a negligible effect on the contrast of the illumination pattern. The beam was then expanded to a diameter of 10 mm using a pair of achromatic doublets before it was incident on a LCoS SLM (SXGA-3DM, Forth Dimension Displays). The SLM was configured to display a series of binary phase gratings, where each grating pixel acts as a switchable half waveplate and imparts a phase delay of either 0 or π ^11^. An image of the SLM was then projected onto the sample via a pair of 4f imaging systems: the first formed by a pair of achromatic doublets and the second by the tube and objective lenses of the microscope. Unwanted diffraction orders, due to the pattern displayed on the SLM and its underlying pixilation, were removed using a custom spatial filter mounted at the Fourier plane of the SLM (the back focal plane of the first achromatic doublet after the SLM). The filter contains three pairs of 0.5 mm diameter holes punched into an aluminium disc at locations equally spaced around the circumference of a circle of diameter either 1.99 mm or 1.49 mm for a nine-or twelve-pixel SLM grating period respectively. The microscope was fitted with a quad band filter cube (TRF89901-EM – ET, Chroma) and fluorescence images were recorded using a scientific CMOS camera (ORCA Flash 4.0, Hamamatsu Photonics).

**Fig. 2.**
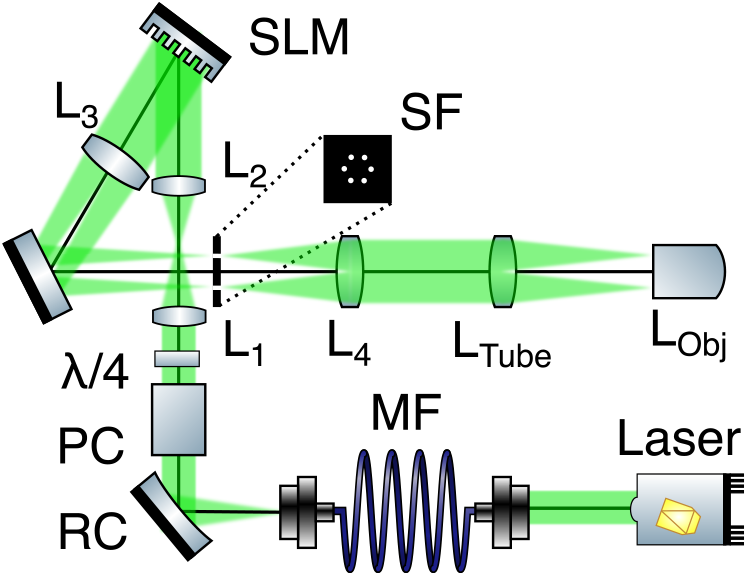
Schematic showing the optical design of the SIM system. Abbreviations: PC : pockels cell; RC: Reflective collimating mirror; MF: multimode fibre; λ/4-achromatic quarter waveplate; L_1_-L_4_ −achromatic doublets; SLM-spatial light modulator; SF: spatial filter for removing unwanted diffracted orders; L_tube_ – tube lens; L_Obj_ -Objective lens.

SIM is especially sensitive to imaging aberrations as the reconstruction algorithms rely on a precise knowledge of both the experimental point spread function (PSF) and the illumination pattern ^19,20^. Our SIM system uses a silicone oil immersion lens (60x/1.3 UPLSAPO60XS, Olympus) to minimise aberrations arising from a refractive index mismatch between the immersion medium and the sample ^21^ (typically cells or tissues in an aqueous buffer or closely matched mounting media). For 3D image reconstruction, using passband attenuation to remove out of focus information ^10^, the SLM was typically programmed to display a series of binary gratings with a period of either nine or twelve pixels (depending on sample thickness and fluorophore brightness). These gratings result in illumination of the sample with sinusoidal irradiance patterns of spatial frequency (period) 3.4 µm^-1^ (294 nm) and 2.6 µm^-1^ (392 nm) respectively, which correspond to 64% and 48% of the incoherent cutoff frequency of the objective lens at 488 nm. Focal series (z-stacks) were captured using a fast-piezoelectric actuator (NanoScanZ, Prior Scientific) to scan the sample axially.

### SLM and camera configuration

A sequence (running order) of illumination patterns was stored in the SLM’s onboard memory as a set of binary images (bitplanes) allowing fast update of displayed gratings. The initialization of the sequence and advance to the next bitplane was controlled using three onboard TTL trigger pins. Specifically, the SLM was programmed to display the first bitplane in the sequence when it received primary HIGH signal. An additional secondary falling edge was then used to trigger the advance to the next frame. Once the primary signal was pulsed LOW, the bitplane sequence was reset to ensure a consistent ordering of the illumination patterns within each raw SIM image sequence of nine images (for an example SLM configuration ‘repertoire’ file see ^22^). The system is typically operated using the same set of nine SLM bitplanes for all excitation lasers which means that the shift of the first order information passbands is the same for all wavelengths, whereas to achieve the theoretically maximal increase in spatial resolution requires a wavelength dependent shift. This simplifies the experimental system, as for all three pattern orientations the gratings are straightforward to generate and can be phase shifted by 2π/3 radians on the pixelated SLM. In practice this does not generally have a significant effect on the quality of the reconstructed images, particularly for closely spaced excitation wavelengths. For applications where it is critical to maximise spatial resolution or when performing SIM using total Internal reflection fluorescence (TIRF) excitation (where the first diffracted orders from the SLM must be maintained within the TIRF ring of the objective lens pupil) the mmSIM framework can be modified to accommodate different bitplanes for different excitation wavelengths. A simple way to achieve this is by creating an extended running order containing all required bitplanes (nine per excitation wavelength) and then modifying the µManager startup script (see ‘Hardware synchronisation’) such that the SLM reset trigger signal is sent after acquisition of all colour channels. The downside of this approach is that a unique running order must be created for each combination of excitation wavelengths. This is relatively simple, for example using the algorithm described in ^14^ to create individual bitplanes.

To match the field of view (FoV) captured by the camera to the size of the illuminated region of the sample, a 512 × 512 pixel windowed ROI close to the centre of the camera sensor was read out for each image giving a maximum theoretical frame rate of 400 Hz or 44.4 super-resolution frames per second. In practice the achievable theoretical frame rate was also limited by the response time of the SLM (∼1 ms). The global exposure period of the camera’s rolling shutter (the interval during which all rows in the sensor are read out) was used to synchronise the acquisition of images with the update of bitplanes displayed on the SLM.

## SIM control software

### Micro-manager hardware configuration

µManager (version 2.0–gamma nightly build) was used to control all microscope hardware. µManager is an open-source project that extends ImageJ ^23^ to control microscopy hardware and comes fully featured with a large library of useful drivers. It provides a convenient GUI interface, several plugins for extending microscopy capabilities and a long list of supported hardware. In-built features and acquisition modes that are particularly use for SIM include multi-dimensional (xyzct) image acquisition, which allows simple multi-colour time lapse imaging and the capture of multiple image fields which can be fused post capture into a single large FoV image. This latter feature is particularly important as SLM-based SIM systems typically have a small field of view due to the limited number of SLM pixels. µManager’s image-based autofocus routines are useful for long duration time lapse imaging, enabling correction of focus drifts without additional hardware.

All the addressable hardware devices (excluding the SLM) were connected to, and interfaced through, µManager using the *Hardware configuration wizard*. The wizard provides an extensive list of available devices under suitable categories; which, once selected, add useful properties of all the devices to the *Device/Property browser*. For laser power control, we added each laser as a separate *Group*, each *Preset* on the main interface window becomes a convenient intensity slider bar (see Fig. 4(c)). To ensure that the lasers failed safe (no emission) under no voltage (which is particularly important for the 405 nm laser which has potential to damage to the SLM), digital/analogue shutters were set-up whereby a voltage signal (analogue or digital) was assigned as a custom shutter in µManager. Custom shutters open and close during acquisition and convert this to a device HIGH/LOW response which is set in the *Device/Property* browser. It is also both possible and useful to mix physical and digital shutters in µManager. For the specific lasers used here it was also possible to use RS232 via USB for shuttering, but due to the additional handshaking this becomes rate limiting, with a signal latency on the order of 20 ms. A fast-switching physical shutter (Prior Scientific), interfaced through the microscope stage controller (ProScan III, Prior Scientific) was added to switch on/off the brightfield illumination in µManager.

All necessary device properties were added as a separate configuration group and the requisite device properties and settings were added to a preset named *Startup*; for safety this group set all laser power levels to zero and all shutters closed upon initialisation. A configuration file for the system is provided as an example. In addition, a µManager startup script (*MMStartup*.*bsh)* ^22^ was used to set the region of interest of the camera to the area of the sample illuminated by the SLM as this cannot be set through the *Device/Property* browser and so could not be saved in the hardware configuration file.

### Hardware synchronisation

Precise timings between the addressable hardware components are essential to ensure the rapid, reliable acquisition of raw SIM image data. µManager was used as the primary source of software triggering signals for the camera, filter wheel and XY and Z stages, and controlled initiation of hardware trigger signals to reset the SLM running order and control the laser shutters. The camera itself acted as a secondary trigger source for advancing bitplanes within the SLM running order. To achieve this a data acquisition (DAQ) card (PCI-6733, National Instruments) was controlled by µManager and used to send TTL voltage pulses to control the camera frame acquisition, the advance of bitplanes in the SLM running order and the switching of laser lines. The Pockels cell was driven by an analogue voltage also produced by the DAQ. Laser illumination intensity was set prior to image acquisition using an analogue voltage signal from the DAQ (see Fig. 3). Using µManager as the primary for timing and synchronisation ensures that each process (such as the switching of laser lines, emission filter and execution of the autofocusing routine) runs to completion before the following process begins.

**Fig. 3.**
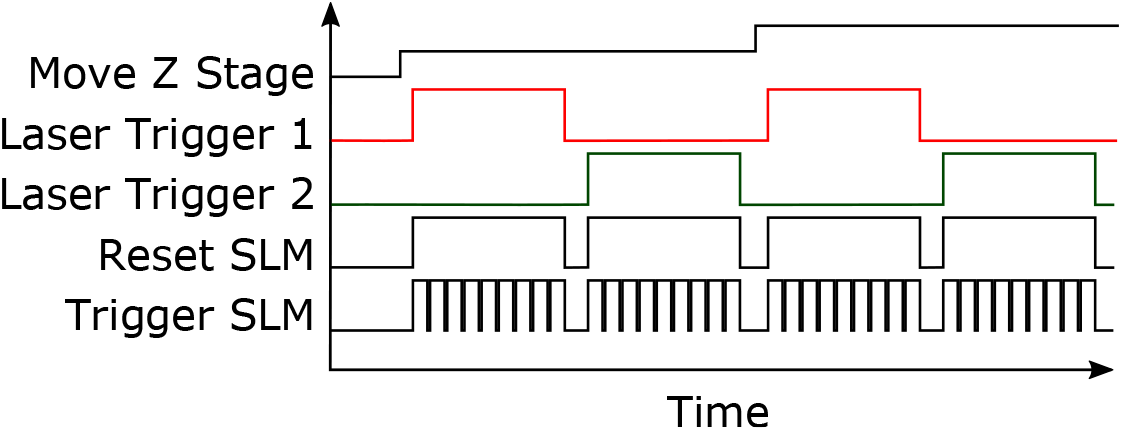
Trigger signals for synchronisation of SIM hardware. Each SIM image requires the capture of a sequence of nine raw images with the camera acquisition synchronised to the display of bitplanes on the SLM. At the beginning of each image sequence the SLM reset signal is set HIGH, causing a reset of the SLM bitplane repertoire at the next falling edge. Colour channels are captured sequentially using a corresponding emission filter. For each colour channel the z stage is moved through a series of predefined position to collect a focal series (z-stack).

**Fig. 4.**
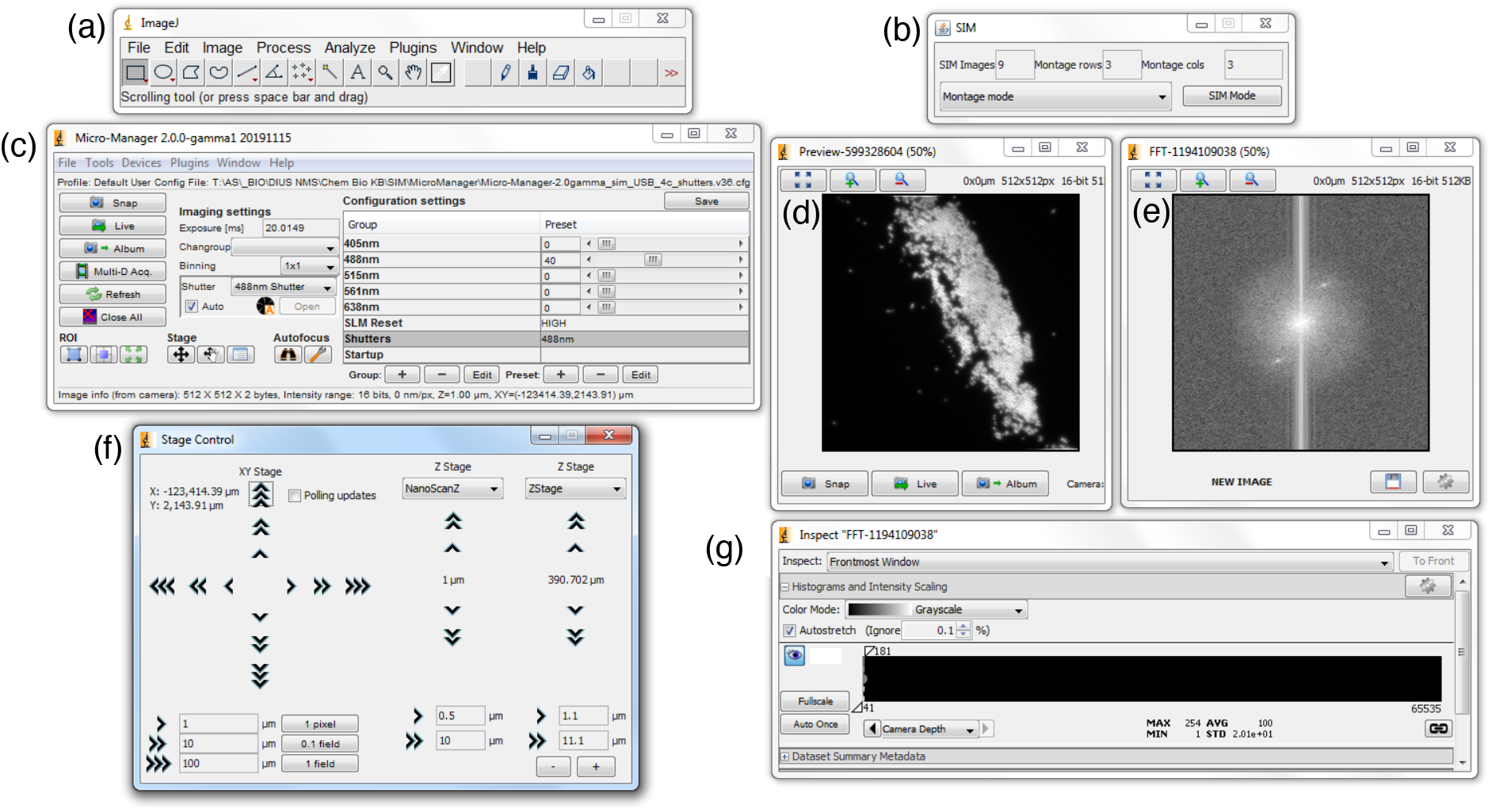
GUI interface for SIM control using Micro-Manager. *SIM Mode (b)* is activated through the GUI forcing Micromanager to produce SIM montages (3 x 3 images containing raw sample images captured under different illumination pattern phases and orientations). The *LiveFFT* window (e) can be accessed from the live preview window (d) (with (g) being the inspection window), the bright spots in the live FFT window correspond to the frequency and orientation of the real space sinusoidal illumination pattern. (c) shows the main Micro-Manager panel for controlling the system with (f) being the XYZ stage controller.

The startup script ^22^ was used to send a TTL pulse from µManager via the DAQ to reset the SLM running order, ensuring a consistent ordering of raw images in each SIM dataset. It is also possible to assign the TTL reset trigger to a µManager custom shutter, as with the digital shutters for the laser lines. It is then possible to assign each laser shutter and the reset shutter to a *Multi-shutter* so that each are on HIGH during an acquisition. The timing and synchronization of devices for an image acquisition sequence is shown in Fig. 3. Firstly, the camera is intialised to collect a sequence of (typically) nine images. The SLM running order is then reset using a HIGH TTL. Concurrently, the emission filter wheel is software triggered to position the correct filter in front of the camera and the XY sample stage is moved (if needed) before the relevant laser shutter is opened using a TTL signal. The camera is then software triggered to begin image acquisition. For each frame collected, the SLM is triggered to advance to the next illumination pattern using the falling edge of the SENSOR_READOUT (end of the global exposure period) trigger signal from the camera. Once a complete sequence of nine images has been collected µManager sends a LOW TTL signal to the SLM which resets the running order. This cycle is then repeated for each SIM image collected in a multi-dimensional acquisition.

## mmSIM plugins

To extend µManager with useful SIM functionality we developed two additional plugins, *mmLiveFFT* and *mmSIMCapture* ^22^. These plugins are easily installed by copying the corresponding .*jar* files directly into the *Micro-Manager-2*.*0gamma\mmplugins* folder.

### Live FFT plugin (mmLiveFFT)

After installation *mmLiveFFT* can be accessed by clicking the cog symbol in the lower right-hand corner of the image preview window. The feature hooks the live preview window and produces a 2D power spectrum (log of the amplitude of the fast Fourier transform) of the current image in a new window (see Fig. 4(e)). We find the spectrum display is useful for checking that the illumination pattern is optimally focused at the front focal plane of the microscope objective – evidenced by two bright points in Fourier space. This is important for daily health-checks on the system as well as being helpful for the alignment of microscope components during construction, maintenance and servicing. The signal-to-noise ratio in the first order information passbands (which carry high spatial frequency information) is strongly dependent on pattern contrast and, as a result, co-alignment of the illumination pattern with the front focal plane of the microscope objective is essential for effective SIM imaging. High pattern contrast is also important for reliable calibration of the illumination pattern parameters. If the estimation of these parameters is inaccurate, computationally inverting the frequency mixing process results in artefacts in the final image. We also find live display of the FFT is useful both for focusing the microscope when imaging thin samples and also in setting the optimal position of the objective lens correction collar. Currently the *mmLiveFFT* plugin is not GPU accelerated but is sufficient^*^ for live update at video rates (20Hz). However, it is advisable to disable the live FFT prior to data acquisition to minimize potential acquisition delays.

### Acquisition plugin (mmSIMCapture)

The SIM acquisition plugin (*mmSIMCapture*) provides a simple GUI (see Fig. 4(b)) for activating *SIMMode* wherein µManager rapidly captures a sequence of nine images for each software triggered acquisition. The resulting images are collected and bundled into a 3×3 image montage to avoid µManager’s (current) restrictions on multi-dimensional image storage ^24^. Once collated the images are displayed and stored in the directory defined by the user, with the suffix *_SIM*. The plugin works by exploiting µManager’s *Runnables* system which provides a programmatic interface to run code every time µManager attempts to acquire an image. The plugin is designed to be non-invasive and as such reverts any settings it changes during execution. This ensures it is safe to run an experiment with the shutter closing between image capture in a *Multi-Dimensional Acquisition*, which is particularly useful for long time-lapse acquisitions. The plugin has been successfully tested for crashes and or memory leaks during a day-long time-lapse experiment. µManager has a built-in event handler for dealing with high-speed imaging and processing, which we find performs reliably under most reasonable imaging conditions.

### Sister SIM System Design

A second SIM system was developed using the same basic hardware architecture again controlled through µManager using the *mmSIM* tools presented here. This second system incorporated two DPSS lasers (Sapphire 488 and Sapphire 561, Coherent Inc.), combined using a dichroic mirror. Both beams were expanded onto the SLM as for the previous system. Unlike the lasers in the previously described SIM system the DPSS lasers could not be rapidly modulated directly and were switched on and off using a physical shutter (SH05/M, Thorlabs Inc.) mounted in the beam path after the dichroic reflector. The microscope used a lower magnification silicone immersion objective lens (30x/1.05 UPLSAPO30XS, Olympus), to provide a larger FoV at the expense of a small reduction in spatial resolution. Images were recorded with a CCD camera (ORCA R2, Hamamatsu). Advance of bitplanes in the SLM running order was again initiated by TTL signals from the camera, but in this case repeated through a microcontroller (Arduino Mega 2560) to improve the fast-switching TTL signal at high frame rates. One limitation of this system stems from the reduced triggering functionality of the camera and the fact that the camera controller continually emitted a heartbeat of HIGH to LOW TTL signals independently of image acquisition. This meant it was not possible to reset the running order of SLM bitplanes at the start of each image acquisition resulting in a variable ordering of raw images in each SIM image sequence. To accommodate this, each image sequence was reordered post capture as a first step in the image reconstruction process.

### Calibration, image reconstruction and testing

Both SIM microscopes are calibrated by imaging a pre-prepared slide of fluorescent microspheres (PS-Speck, Thermo Fisher Scientific) deposited onto a (#1.5H) coverslip. After air drying, a drop of hard curing mountant (ProLong Gold, Thermo Fisher Scientific) was added and a glass microscope slide was placed on top. The monolayers of closely packed microspheres which form on parts of the slide are ideal for calibrating the illumination pattern parameters (frequency and orientation of the sinusoidal illumination patterns). We use in-house developed MATLAB routines for calibration of the illumination pattern and reconstruction of super-resolution images based on the method described in ^9^. Briefly, illumination parameters are estimated by cross correlation of separated zero and first order Fourier space passbands. Phase offsets between the passbands are determined by complex linear regression in overlapping areas after accounting for OTF differences. Separated passbands are then multiplied by an inverted Gaussian to attenuate out of focus information ^11^, shifted in Fourier space (by multiplication of the real space image with the corresponding cosine) and added together. Once the process has been repeated for three pattern orientations an inverse Fourier transform yields the final high-resolution image. A good alternative software package, based on the same principles, is the open source fairSIM program ^15^ which is freely available through FIJI. It is recommended that all reconstructed images are run through SIMcheck ^14^ in order to observe and diagnose potential artefacts.

The speed of the primary SIM system was tested by imaging the above calibration slide. The minimum camera exposure time for which we were able to acquire and reconstruct SIM images was 5.5 ms, corresponding to 49.5 ms for capture of the nine raw images in a SIM image sequence and a super-resolution frame rate of 20 frames per second (fps). For shorter exposure times hardware synchronization and triggering became unreliable. In practice the achievable frame rate is limited by the brightness of the fluorescent sample, the photostability of the fluorophores and the tolerance of the specimen to phototoxic effects resulting from high irradiance levels.

Imaging performance of the microscope system was assessed by imaging a slide containing conjugated polymer nanoparticles (CPN550, Stream Bio) prepared using the same method as described previously for the calibration slide. CPNs are fluorescent nanoparticles comprising an encapsulated semiconductor light emitting polymer core and have a nominal diameter of 70 nm. The split view image shown in Fig. 5(a) shows a clear reduction in the apparent size of the nanoparticles due to the improvement of lateral spatial resolution in the SIM image. This is accompanied by a significant increase in image contrast which enables visualization of individual nanoparticles within the densely packed monolayer. As a second test of the system, we imaged A549 (human lung carcinoma) cells stained for mitochondria (MitoTracker Deep Red, Thermo Fisher Scientific). The split image (Fig 5. (b)) shows maximum intensity projections of SIM and corresponding diffraction limited widefield z-stacks through a single cell. The dramatic improvement in both spatial resolution and image contrast in the SIM image allows individual isolated mitochrondria and fine filaments to be clearly identified.

**Fig. 5.**
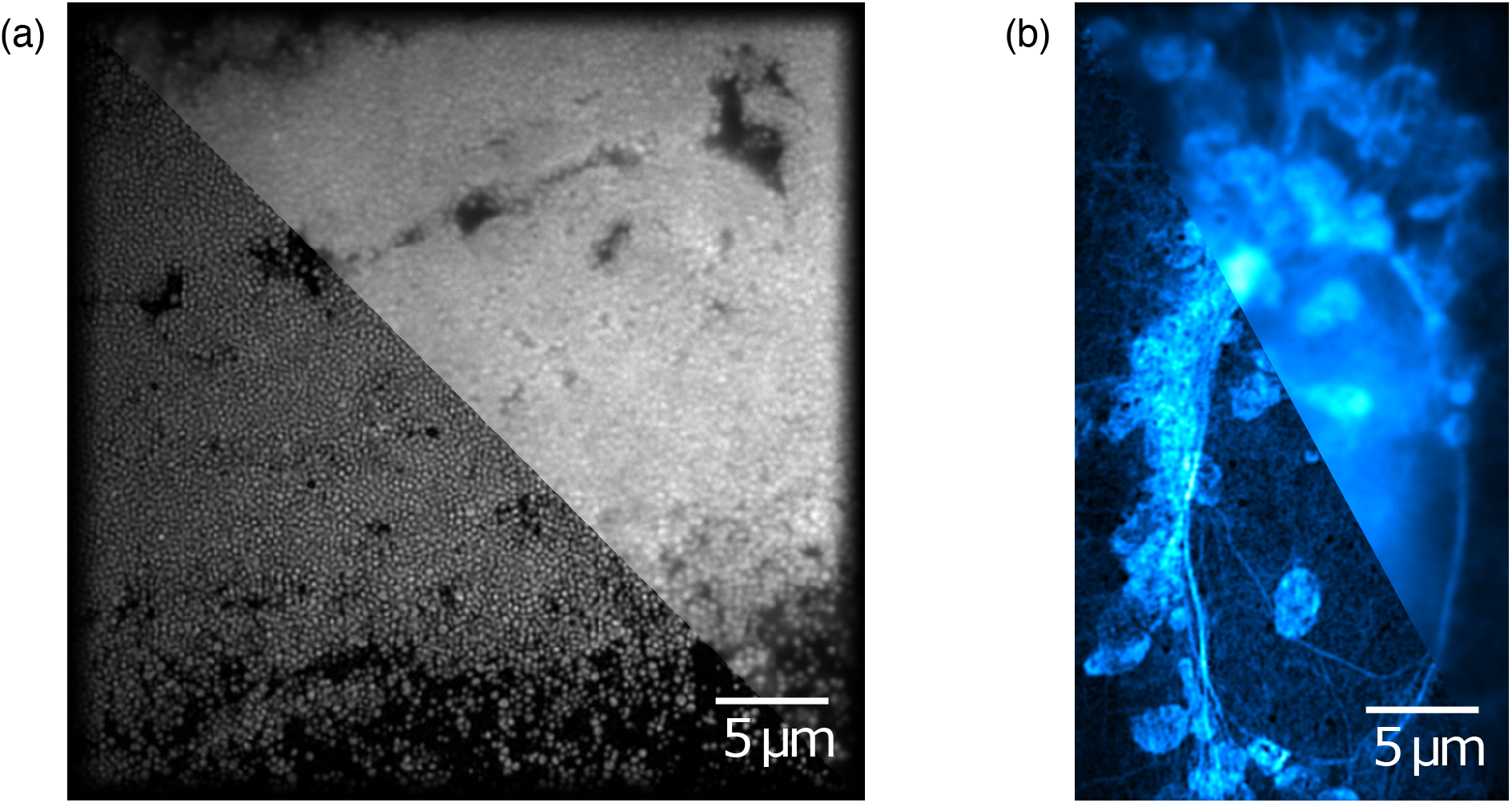
Split images showing diffraction limited (upper right) and corresponding reconstructed SIM image (lower left). (a) Yellow conjugated polymer nanoparticles dried onto a glass coverslip. (b) A549 cell stained for mitochondria (MitoTracker Deep Red).

## Conclusions

This article describes a new approach to SIM hardware control and image acquisition based on the open source µManager package. We provide instructions for configuration, control and synchronization of SIM system hardware and introduce two new plugins (*mmLiveFFT*) and (*mmSIMCapture*) to allow real time visualization and simple capture of raw SIM image data. These new software tools have been used to develop two home-built systems using widely available off-the-shelf components. The resulting microscopes are capable of fast (up to 20 fps), robust, high quality, optically sectioned super-resolution imaging. Combined with the large number of in-built µManager features, this approach allows the simple development of sophisticated programmatic, multi-dimensional imaging workflows important for a wide range of bioimaging applications. By removing a significant barrier to developing a flexible, customisable home-built SIM we believe our work has the potential to enable further innovations in SIM. Finally, the methods and tools presented here for extending µManager may also prove useful for establishing microscope systems based on other super-resolution techniques, such as single molecule localisation microscopy, which trade off temporal resolution for spatial resolution.

## Additional Information

## Acknowledgments

The authors thank Nilofar Faruqui from the National Physical Laboratory for preparing MitoTracker labelled A549 cells. Conjugated polymer nanoparticles were provided by Dermott O’Callaghan from Stream Bio and prepared for imaging by Camilla Dondi from the National Physical Laboratory.

## Data Accessibility

All source code, documentation and files are available through GitHub (https://github.com/ctr26/MMSIM) and archived using Zenodo (https://doi.org/10.5281/zenodo.4432274) ^22^.

## Author Contributions

CTR designed and developed the mmSIM software and implemented it on microscope hardware designed and built by MJS. MJS benchmarked the performance of the software and collected imaging data. CTR and MJS wrote the manuscript.

## Footnotes

* Tested on a Dell Optiplex (3.3 Hz quad core CPU (Intel Xeon E5-2643); 48 GB RAM running Windows 7) and a 2019 Macbook Pro

